# *Drosophila* model of Anti-retroviral therapy Induced Peripheral Neuropathy and Nociceptive Hypersensitivity

**DOI:** 10.1101/2020.06.18.160077

**Authors:** Keegan M. Bush, Kara R. Barber, Jade A. Martinez, Shao-Jun Tang, Yogesh P. Wairkar

**Affiliations:** Neuroscience Graduate Program, University of. Texas Medical Branch, Galveston, TX 77555; Mitchell Center for Neurodegenerative Diseases, Department of Neurology, University of Texas Medical Branch, TX 77555; Department of Neuroscience, Cell Biology, and Anatomy, University of Texas Medical Branch, TX 77555

## Abstract

The success of antiretroviral therapy (ART) has improved the survival of HIV-infected patients significantly. However, significant numbers of patients on ART whose HIV disease is well controlled show peripheral sensory neuropathy (PSN), suggesting that ART may cause PSN. Although the nucleoside reverse transcriptase inhibitors (NRTIs), one of the vital components of ART, are thought to contribute to PSN, the mechanisms underlying the PSN induced by NRTIs are unclear. In this study, we developed a *Drosophila* model of NRTI-induced PSN that recapitulates the salient features observed in patients undergoing ART: PSN and nociceptive hypersensitivity. Furthermore, our data demonstrate that pathways known to suppress PSN induced by chemotherapeutic drugs are ineffective in suppressing the PSN or nociception induced by NRTIs. Instead, we found that increased dynamics of a peripheral sensory neuron may underlie NRTI-induced PSN and nociception. Our model provides a solid platform in which to investigate further mechanisms of ART-induced PSN and nociceptive hypersensitivity.

## Introduction

While the antiretroviral therapy (ART) has been successful in controlling the symptoms in acquired immunodeficiency syndrome (AIDS) patients, a significant proportion of patients on ART develop peripheral sensory neuropathy (PSN)^1–7^. These symptoms include pain and/ or numbness in the extremities^8^. Thus, while NRTIs can control the HIV infection, they can also cause neurological complications^11–16^. Therefore, chronic administration of ART may also contribute to the neurological complications, including PSN ^13,14,17–20^. In particular, nucleoside reverse transcriptase inhibitors (NRTI), the backbone of ART regimens, are especially relevant in this regard because their neurotoxicity is well established^17,21–25^. For example, d4T, an NRTI still used in some resource-limited countries, has shown to be neurotoxic not only in HIV-patients but even in uninfected subjects that received it as prophylaxis^26^. However, little is known about the mechanisms underlying the neurotoxicity of NRTIs in peripheral nerves. Previous studies have suggested various mechanisms of neurotoxicity, including mitochondrial damage ^27–29^ and neuroinflammation ^24,30,31^. Thus, the current research on the neurotoxicity of NRTIs is focused on describing their detrimental impacts on specific biological pathways such as mitochondrial homeostasis and neuronal apoptosis. While these approaches have been helpful, the mechanistic insights into the effects of NTRI-regulated biological pathways to the PSN have not been conclusively established. We have taken the advantage of the similarity between *Drosophila* and mammalian sensory neurons ^32–34^ to establish a novel strategy that will allow for genome-wide unbiased forward genetic screens to understand the molecular mechanisms that play key roles in regulating the development of NRTI-induced neurotoxicity in the peripheral nerves. Interestingly, when *Drosophila* larvae are subjected to NRTI treatment, the peripheral branches of sensory neuron dendrites show an increased instability and clear signs of fragmentation as compared to the untreated larvae. Moreover, genetically restoring stability to the dendrites of the peripheral sensory neurons significantly suppresses their degeneration. In addition to the sensory neuron fragmentation, the larvae where the sensory neurons are genetically stabilized also show a significant reduction in nociceptive hypersensitivity, indicating that the instability of peripheral sensory neurons might drive the degeneration and the nociceptive hypersensitivity in the *Drosophila* model. Thus, our study provides a genetically amenable platform to further dissect the molecular pathways underlying NRTI-induced PSN and nociceptive hypersensitivity.

## Methods

### General methods

All the flies were cultured at room temperatures. All the crosses were also performed at the room temperature. In all experiments involving nociception assays, adult *Drosophila* males and cirgin females were transferred to food containing vehicle, which usually was water, except DMSO that was used for experiments with pertaining to Taxol. In all experiments where larvae were utilized, only wandering third instar larva were used. All experiments within a figure panel were performed on the same day by the same experimenter. Experiments where the expression in C4da neurons was utilized, the control flies were ppk-EGFP5. Otherwise, Canton S. was used as a control unless specifically stated. For each cross utilized throughout the study, ~10 virgin females were crossed to 7-8 males. These adult flies were allowed to lay eggs for 2-5 days per vial and then transferred out of the vial. Therefore, in all experiments, the larvae developed in the food containing the drug or no drug.

### Thermal Nociception Assay

Wandering third instar larva were placed in ~1mL of water in individual wells of a 96 well plate. If more than one genotype was being tested, they were placed opposite their drugged counterparts so that the wells were equidistant from the center of the block to ensure equal heat distribution. A black thermally conductive backplate was placed beneath the well plate for contrast so that the camera could track the larval movements with ease. A sensitive temperature probe was submerged in a centrally located unoccupied well with an identical volume of water. A camera was placed such that the wells with larvae and the temperature probe readout were clearly within the view of the camera. The recording of thermal nociception was started while the hot plate was at room temperature. The initial set point of the hot plate was RT and the temperature was raised in 0.1°C/10 seconds increments. This rate was maintained throughout until the temperature reached 40°C. Videos were recorded for the entire period and the videos were quantified by two independent experimenters for the larval writhing response.

The nociceptive behavior used to qualify as nociceptive writhe is a well-documented corkscrew response^32^. The thermal nociception temperature at which each larva exhibited the corkscrew-like writhe response was recorded from the temperature of the probe inside the water bath. At 40°C, most of the larva died and therefore, this last set of readings were not included in analyses. After analyzing the videos, 3-continuous corkscrew-like rolls were scored as nociceptive writhe. This temperature was designated as the minimum temperature required for the writhing response (Movie 1, 2).

### Heat Probe Nociception Assay and Quantification

Heat probe nociception assay was performed as described previously ^36^. Wandering third instar larvae raised at 22°C were collected and rinsed in PBS. Individual larvae were tested by touching the heat probe consistently to the posterior third of the larvae. All the experiments were performed at the same time by the same experimenter. The larval response was recorded as non-responding, slow responding, and fast responding groups based on the following criteria: <u>Non-responders</u>-larvae that did not exhibit nociceptive writhe within 20 seconds of the application of the thermal probe. <u>Slow-responders</u>-Larvae that exhibited nociceptive writhe between 5 and 20 seconds of the application of the thermal probe and <u>Fast-responders</u>-larvae that exhibited nociceptive writhe as soon as the probe was applied or within 5 seconds of the application of the thermal probe.

### Mechanical Nociception

Von Frey filaments were designed such that they had similar diameter and flat tips. This protocol was adapted from Kim et al., ^32,37^. These flat tips exert increasing pressure rather than force. The later being the one used to test nociception in vertebrate animals. The filaments used in this paper were made in-house using a series of 12 calibrated monofilament fishing line segments (Berkley Trilene XL Monofilament Fishing Line), which were tethered to a hard-plastic handle. The filaments were calibrated by their ability to depress a balance consistently when vertically exerting a force until bent at roughly 30° from tip to tip. The series of 12 filaments consistently exuded following pressures: (100kPa (0.50mN), 150kPa (0.75mN), 200kPa (1.01mN), 250kPa (1.26mN), 300kPa (2.36mN), 500kPa (3.93mN), 700kPa (5.50mN), 1000kPa (7.85mN), 2000kPa (15.71mN), 3500kPa (27.49mN), 5000kPa (39.27mN)) over a 0.1mm inserted tungsten wire tip. Experiments were performed beginning with the lowest pressure filament of 100kPa (0.50mN). Single larva was secured loosely in place with forceps and the filament tip was consistently pressed straight down on the lower third of the dorsal surface (preferably segment A6) of the larvae and released quickly. The larva was then observed for 30 seconds for nociceptive behavior. If no noticeable nociceptive behavior was observed, the same larva was secured again and tested with the next filament in a sequence until a filament elicited the nociceptive behavior (writhe). We held the temperature of the test environment constant at 25-27°C because in our hands it affected the sensitivity of the larvae. Notably, temperatures below 24°C significantly decreased the larval response.

### Larval Motility Assay

Larval motility was assessed based on the method described in Nichols et al, 2012 ^38^. Wandering third instar larvae raised at 22°C were collected and rinsed in PBS. Individual larvae were assessed for their locomotor activity utilizing a plastic-covered graph paper with 1cm^2^ grid lines. Larvae were allowed to move freely for 1 minute and the number of lines crossed was noted.

### *Drosophila* NRTI treatment

All flies were raised at room temperature (22°C) with natural day night cycle. WT (*Canton S*) flies were placed in vials containing 2.5ml of instant *Drosophila* media (“blue food”, Carolina Biological Supply, Burlington, NC) made with water base. NRTI experiments utilized water as a vehicle and the additional water in each NRTI vial amounts to less than 1μL. NRTI dose response concentrations for **AZT** are: (0.026 μg/mL, 0.13 μg/mL, 0.26μg/mL (main concentration used throughout), 1.3 μg/mL, 2.6 μg/mL, 13 μg/mL, 26 μg/mL, 52 μg/mL, 104 μg/mL, 208 μg/mL, and 416 μg/mL); or **ddC** (0.0014 μg/mL, 0.014 μg/mL, 0.07 μg/mL, 0.14 μg/mL, 0.28 μg/mL, 0.56 μg/mL, 0.84 μg/mL, 1.12 μg/mL, 1.4 μg/mL, and 2.8 μg/mL). The AZT and ddC are water-soluble and were stored in 1 mg/ml and 3.3 mg/ml concentrations respectively. AZT (Catalog# 3485) and ddC (Catalog # 220) were obtained from the NIH AIDS Reagent program (http://aidsreagent.org). Taxol experiments were treated with DMSO (Vehicle), with or without 30μM of 1 mg/ml Taxol (Paclitaxel, Sigma, St. Louis, MO) in DMSO. A stock of 1 mg/ml paclitaxel in DMSO was utilized for dilutions with identical amounts of DMSO used as control.

ppk-EGFP5 virgin female flies (10) were crossed to males (7-8) required for the particular experiment in water-based food (Control) and food treated with 0.26μg/mL AZT diluted in similar amounts of water. Every cross in the study was made using at least 10 virgin females and 7-8 males. These adult flies were allowed to lay eggs for 3-5 days per vial. These crosses were maintained at room temperature (22°C). No significant differences were noted between males and female larvae or adult flies and their response towards the nociceptive stimuli, therefore, the data was collated for analyses.

### Dissection, Imaging, and Analyses

Specific imaging of C4da neurons was achieved via live imaging of ppk-EGFP5 lines that have been well-characterized^63^. No immunostaining was used. All images were acquired using a Nikon Eclipse 90i laser scanning confocal microscope with either a 20X air or 60X oil objective. C4da neurons were visualized using the EGFP fluorescence of ppk-**EGFP5**. Each larva was mounted individually for imaging. Where the larvae were dissected, the dissections were performed in the following way: Larvae were pinned at head and tail submerged in cold HL3 solution. An incision was made horizontally across the larval cuticle near the tail pin in such a way that only one side of the midline from pin to pin was cut. A final incision was then made near the headpin. This method helped maintain total integrity of one-half of the Class IV da neurons (C4da), which was used for analyses. In our hands, the integrity of this one side of the dendrites was maintained for as long as 15 minutes. This was confirmed by imaging the dendrites to find discontinuities of staining in GFP. In all cases, the staining appeared smooth and uniform throughout the C4da neurons. The trachea and guts were carefully removed using forceps and the larvae were unpinned and removed from the dissection plate to be placed cuticle side up on a glass slide. A small drop of cold HL3 was added to the larva on the slide before a small square coverslip was placed on top carefully to flatten and maintain the larval position for immediate imaging. The time taken from the first incision to the start of imaging was recorded and maintained within 2 minutes of variation; which for the entire experiment did not exceed 5 minutes (From the first cut to the imaging). Single C4da neuron and its dendritic field were imaged per larva from abdominal segment 3 or 4 with a laser dwell time of 1.68 μs per pixel at 1024 × 1024 spatial resolution. The sensory neuron was imaged through its entire z-dimension with a step size of 1 μm. The exception to this imaging method is the experiments in Figure 6 where Canton S. and *Ect4*^−/−^ *Drosophila* lines were utilized. In this particular instance, the images shown are immunofluorescent labeled neuronal membrane marker, HRP. Therefore, this figure does not show specific labeling of C4da neurons but instead labels all the sensory neurons. Apart from the fluorescence labeling, everything else was kept identical to the method described above.

For live imaging, larvae were rinsed in PBS and were directly mounted (without dissecting) within a coverslip cage in the HL3 solution. The coverslip cage was made to house the larva under slight pressure to hold it in place during the ~3hour imaging period. The cage used in these experiments was made with two coverslips that touched the larvae such that the larval body was in between the coverslips. The dorsal projections of one C4da neuron per larva from abdominal segment 3 were imaged every 10 minutes for 3 hours through the cuticle with a dwell time of 1.68 μs per pixel at 1024 × 1024 spatial resolution with a step size of 2 μm. Approximately, 10μL of fresh HL3 was added to the coverslip cage every 10-15 mins throughout the live imaging process to keep the larvae hydrated and to prevent hypoxia. The motility of larvae was assessed after the live imaging session was completed to assess their general health and only the images from larvae that showed normal motility were used in the analyses.

For analyses of dendrite fragmentation, confocal image stacks were converted to maximum intensity projections using ImageJ software. All analyses were performed on the entire C4da arbors of the single C4da neuron that was imaged. The total numbers of terminal dendritic branches and the total number of fragmented dendritic branches were counted manually, and the count was tracked using ImageJ. Discontinuous GFP staining pattern was analyzed manually and was counted as fragmented terminal dendrite. For analyzing the time-lapse images, confocal images in the time series were converted to maximum intensity projections using ImageJ software, and a random box of an area of 100 × 100 μm was drawn away from the cell bodies. Between each 10min time point image, the total numbers of growing (from the same main branch), retracting, and sprouting (new branch formation) branches were counted manually for the entirety of the 3-hour time-lapse. The minimum length for qualifying as sprouting, growing, or retracting branch, the event had to be >1-2μm in subsequently acquired images.

For determining muscle size, wandering third instar larva were dissected and fixed with Bouin’s fixative, which stains the muscles bright yellow. Bright-field images were taken, and ImageJ was used to quantify the muscle #4 area of abdominal segment 3. At least 10 larvae for each experiment and genotype were analyzed.

### Statistical Analyses

All the experiments were performed with the experimenter blinded to the genotype of the larvae. The coded groups were then subjected to statistics before revealing their identity.

Statistical analyses were performed using the GraphPad Prism software. The Student’s T-test was used for comparison between two groups and one-way ANOVA followed by Bonferroni posthoc test was utilized when comparing multiple groups. Significance levels were set to 95% confidence intervals and the p values are indicated in the respective figure legends.

## Results

### Exposure to *AZT* induces thermal and mechanosensory nociceptive hypersensitivity in *Drosophila*

*Drosophila* larval model has been previously used in understanding the mechanisms of nociception ^32,33,43–45^. When subjected to noxious stimuli, like high temperatures, the larvae respond by a characteristic “corkscrew-like” escape behavior, also called as writhe ^46^, which has been successfully exploited to screen for genes involved in nociception ^34,47,48^. Larvae that are sensitive to these noxious stimuli generally respond with writhe at a lower threshold than the control larvae. We adopted this established behavioral paradigm to test whether exposure to NRTIs can induce nociceptive hypersensitivity in wild-type (WT) larvae. We used a water bath made of polypropylene fitted with a sensitive temperature-measuring probe that can detect temperature fluctuations of 0.1°C (Fig. 1A and Supplementary Movie 1). To test for nociceptive hypersensitivity, the temperature of the water bath was ramped up gradually in 0.1°C/10 increments. A camera attached to the microscope tracked both the rise in temperature and larval movements (Fig. 1A). A writhing response by the larvae was recorded as a nociceptive hypersensitive response if the larvae showed at least three corkscrew-like movements without a stop at a temperature that was lower than the one that induced a similar reaction in WT larvae. First, we sought to optimize the dosage of NRTIs for *Drosophila* larvae. For this, we used a human equivalent dose of two NRTIs: AZT (Zidovudine or Azidothymidine) and ddC (Zalcitabine). Using a recent study that has used drugs mixed in the food to feed larvae^49^, we estimated that 26μg/mL food volume of AZT and 0.14 μg/mL food volume of ddC would be an ideal starting point (see Materials and methods for details). Although this dose induced thermal hypersensitivity in the larvae it also induced a significant amount of lethality (30% in AZT and >80% in ddC, n=20, p<0.01). Therefore, we used two dilutions of this concentration (0.1x, and 0.05X) to establish an optimal dose that would allow us to test nociceptive hypersensitivity without causing significant lethality.

**Figure 1:**
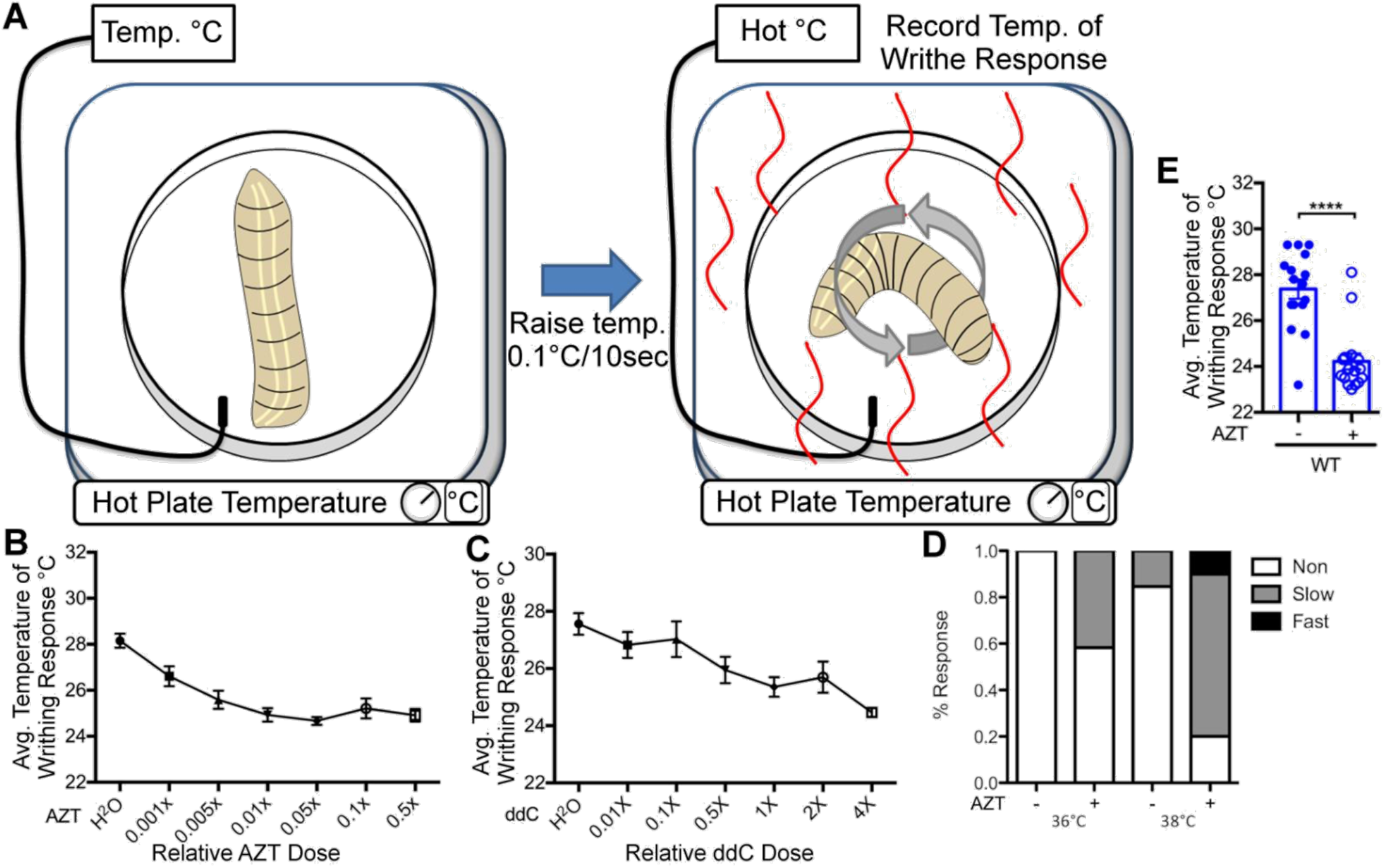
NRTIs induce nociceptive hypersensitivity in *Drosophila*. **A**) Experimental set up for larval thermal nociception assay. Larvae were subjected to small increases in temperatures and monitored for writhing response. The temperature of the writhing response was recorded using a sensitive probe. **B**) Quantification of thermal nociception response of larvae to increasing doses of AZT, dots represent mean and the error bars represent SEM. A total of 16 larvae were utilized per dose with three repetitions of each experiment. F(6, 105) = 14.3, p = 7.68E-12, 1-way ANOVA. **C**) Quantification of writhing response to increasing doses of ddC, dots represent mean and the error bars represent SEM. 9 larva were utilized per dose with three repetitions of each experiment. F(6, 49) = 6.78, p = 2.82E-5, 1-way ANOVA. **D**) Quantification of nociception temperature for WT larva in AZT− and 0.26μg/mL (0.01X) AZT containing food. p = 2.18E-6, T-test. **E**) Quantification for the thermal nociception response to thermal probe at 36°C and 38°C. Larvae raised on AZT exhibit significantly more nociceptive responses at both temperatures. Slow response: larvae responding between 15-20 seconds; Fast response: larvae responding immediately to 5 seconds after thermal probe application.

WT flies were transferred into vials that had drugs (AZT or ddC) mixed into the food. Control flies were transferred on the same day at the exact same time onto food prepared identically, except for the addition of drugs (AZT−/ddC−). Flies were allowed to lay eggs in AZT, ddC, or AZT−/ddC− food for about 3 days before being discarded. Wandering third instar larvae were collected and used for the nociceptive hypersensitivity experiments. We found that larvae raised on AZT or ddC food showed significantly lower temperature thresholds for the nociceptive writhe (at least 3°C) as compared to the control flies raised on AZT−/ddC− food (Fig. 1B, C, and Supplementary Movie 2). This is a significant change given that the thermosensory neurons in flies can sense minute temperature differences ^50,51^. The minimum concentration of drug needed to decrease the temperature to see an observable writhing response with AZT was only 1/100^th^ the estimated dosage at 0.26 μg/mL and did not induce lethality in the larvae (Fig. Supplementary Fig. 1). However, even the minimum concentration of ddC (0.14 μg/mL) required to induce a significant nociceptive hypersensitivity was toxic (50% lethality, n=20, P<0.0001) making AZT a preferred drug to test the effects of NRTI induced neuropathy.

To confirm our data, we utilized another method used to test thermal nociception-the thermal probe method ^36,52^. This method relies on the consistently observable writhing response of larvae when the larvae are touched using a heated probe. This method has two advantages: First, it involves localized heat application as opposed to the global temperature increases encountered by larvae in a water bath. Second, this method allows one to reliably test rapid response to thermal stimulation as opposed to the slow response generated using a temperature bath. When a thermal probe heated to 36°C was touched to the WT larvae raised in AZT−/ddC−, they did not show a significant response. However, the AZT/ddC^−^ larvae start showing a writhing response when the probe is heated to 38°C. Interestingly, 40% of larvae raised on AZT containing food showed a writhing response even at 36°C (Fig. 1D). While 15% of AZT−/dd− larvae also showed a writhing response at 38°C, this response was exaggerated in (80%) larvae raised on AZT at 38°C. Moreover, 15% of them showed a very fast response (Fig. 1D). While these data demonstrate that exposure to AZT induces nociceptive hypersensitivity, we wanted to test whether these responses were driven by the C4da neurons that are responsible for heat sensitivity in *Drosophila* ^52,53^. To test this, we expressed the tetanus toxin light chain (UAS-TeTxLC) in C4da neurons using ppk-Gal4, which specifically silences these neurons^54^. As expected, flies expressing TeTxLC ^53^ showed no response to temperature changes in either AZT/ddC^−^ larvae or larvae raised on AZT, indicating that C4da neurons largely drive the thermal nociceptive hypersensitivity response of NRTIs (Supplementary Figure 2). Finally, as newer NRTIs are introduced regularly, we wanted to test whether these newer NRTIs also induce nociceptive hypersensitivity. Therefore, we performed the same assays with newer NRTIs-Emtricitabine (FTC), Abacavir (ABC), and Tenofovir (Tenovir) (Supplementary Fig 3A). All the newer NRTIs tested showed increased nociceptive hypersensitivity to thermal stimulation, indicating that most NRTIs induce nociceptive hypersensitivity in the *Drosophila* model.

Since anti-retroviral therapy can also lead to the development of mechanical allodynia ^31,57^, we asked whether the larvae exposed to NRTI also showed nociceptive hypersensitivity to mechanical stimuli. To perform these assays, we designed and calibrated Von Frey filaments, in house. Von Frey filaments were calibrated for specific pressures (described in materials and methods) and consistently applied to the posterior third of the larvae (Fig. 2A). Von Frey filaments induced nociceptive writhe in larvae raised on AZT at lower pressures compared to larvae raised on AZT− food, suggesting that exposure to AZT also lowers the threshold to mechanical stimulation (Fig. 2B). Similar results were also obtained using newer NRTIs (Supplementary Fig. 3B). As a control, we also tested whether there were any issues with general motility in larvae raised on NRTIs using the locomotor assay ^38^ and did not find any significant defects in the motility of the larvae (Supplementary Fig. 4A). We also tested for any defects in the development of the larval musculature. We found that the size of the muscles did not differ significantly between the larvae raised on AZT/ddC^−^ food and those raised on AZT (Supplementary Fig. 4B). Finally, the larvae raised on AZT eclosed from pupa at that same time as the WT larvae raised in AZT− food (Data not shown). Together, these data demonstrate that larvae exposed to NRTIs show both thermal and mechanical nociceptive hypersensitivity.

**Figure 2:**
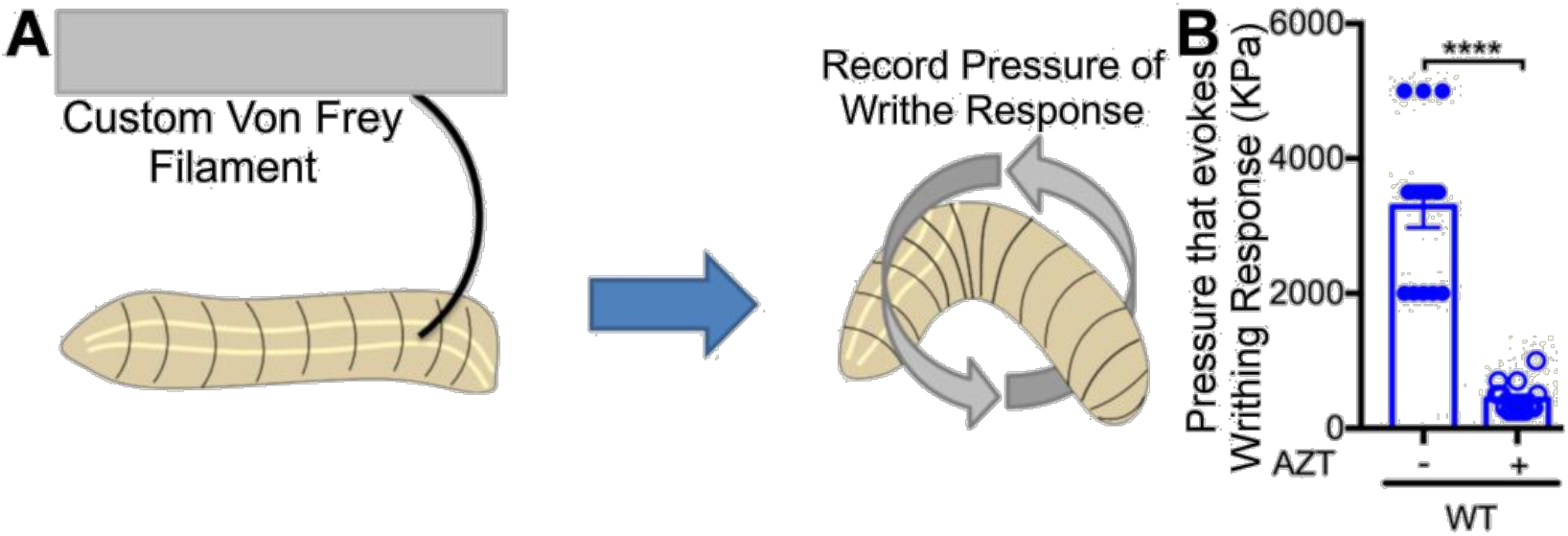
AZT also induces mechanical allodynia. **A**) Diagram of the Von Frey mechanical nociception test for *Drosophila* larva. A set of calibrated Von Frey filaments with increasing pressure from 100kPa – 5000kPa were designed and used to stimulate the dorsal A6 or A7 segment of the larva. Larvae were monitored for 30 seconds for nociceptive behavior before continuing to the next filament. **B**) Quantification of sensitivity to mechanical stimulation in WT larva in AZT− and 0.26μg/mL (0.01X) AZT containing food. p = 1.12E-7, T-test.

### Exposure to AZT shows signs of degeneration in sensory neurons

Skin biopsies of patients with painful neuropathy (Or distal sensory neuropathy, DSP) show degeneration of peripheral sensory neurons^58–61^. To test whether the *Drosophila* model shows signs of peripheral sensory neuron degeneration, we assessed the sensory neurons in larvae exposed to AZT. To test this, we labeled the class 4da (C4da) sensory neurons by driving the expression of EGFP using the C4da neuron-specific Gal4 driver (ppk-Gal4)^63^. These neurons are necessary for nociception induced by noxious temperature and mechanical stimuli ^47,53,63^. Quantification of C4da neuron terminal dendrite branch number and proportion of branches that exhibited fragmentation (Discontinuous GFP Flourescence) revealed that the most obvious difference observed between larvae raised on AZT− media versus those raised on AZT had a significant increase in the proportion of fragmented terminal (distal) dendrites (Fig. 3C), consistent with a decrease in nerve fiber density observed in HIV neuropathies ^58^. Notably, larvae raised on AZT did not have a significant change in the number of terminal dendrites (Fig. 3B), or the primary and the secondary branches of the sensory neurons (Fig. 3D, E). Also, the primary and secondary branches did not show fragmentation phenotype (Fig. 3F). Finally, FTC-a newer NRTI also showed fragmentation similar to that of AZT (Supplementary Fig 5 A, B). Next, we tested whether the terminal dendrite fragmentation caused by AZT was similar to the one that has been reported for paclitaxel, a chemotherapeutic drug ^49^. Consistent with the previous report, we found a significant fragmentation of terminal dendrites in larvae exposed to paclitaxel. However, notably, the fragmentation induced by paclitaxel was much more severe than that induced by AZT (Supplementary Fig. 6). These data indicate that larvae exposed to NRTIs have a significant fragmentation in the terminal branches of sensory neurons.

**Figure 3:**
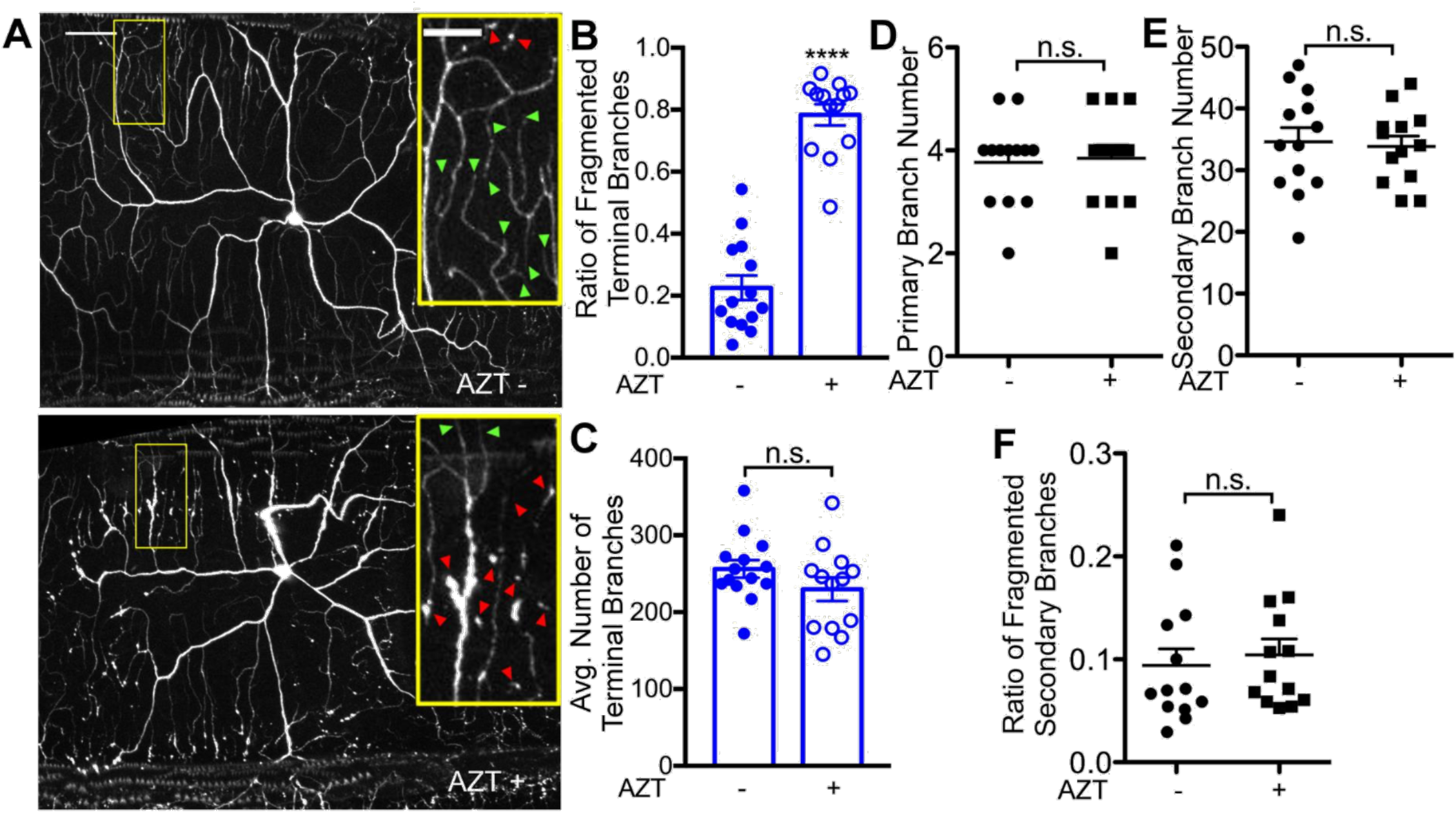
Exposure to AZT leads to the fragmentation of distal dendrites. **A**) Representative images of the C4da sensory neurons of third instar larvae raised on AZT− and AZT+ food. Inset arrows identify terminal branches that are intact (green) and fragmented (red). **B**) The number of terminal branches of C4da sensory neurons is not significantly altered by AZT. p = 0.177, t-test. **C**) The proportion of C4da terminal branches that exhibit fragmentation is increased larvae raised on AZT. p = 9.62E-11, t-test. **D**) The number of primary branches of C4da sensory neurons is not significantly altered by AZT. p = 0.823, t-test. There was no detectable fragmentation of C4da primary branches in AZT− or AZT+ larva. **E**) The number of secondary branches of C4da sensory neurons is unaffected by AZT. p = 0.788, t-test. **F**) The proportion of C4da secondary branches that exhibit fragmentation is unaffected by AZT exposure. p = 0.645, t-test. Also, refer to Supplementary Fig. 6 for C4da neuron response to Taxol. Vehicle (−); AZT +. ****p < 0.0001; error bars = SEM; scale bar = 50 μm; inset scale bar = 20 μm.

### Fragmentation of the terminal dendrites is independent of *wnd*/dlk pathway

The data that both AZT and paclitaxel cause fragmentation of terminal dendrites suggested to us that the mechanisms that underlie the fragmentation might be similar. Work in *Drosophila* and cultured DRG neurons has shown that paclitaxel-induced degeneration can be suppressed by downregulating the Wallenda/di-leucine zipper kinase (DLK) pathway ^49,64^. Interestingly, downregulating DLK also delays Wallerian degeneration-a form of axon degeneration caused due to the damage to axons ^64–67^, suggesting that the DLK pathway might be one of the commons pathways that mediate the injury response signaling.

To test this hypothesis, well-characterized RNAi lines of *wnd*/*dlk* were exposed to AZT along with WT controls ^65,68,69^. If knockdown of *wnd*/dlk blocked the neurotoxicity induced by AZT, we expected to observe a suppression of fragmentation in *wnd*/dlk knockdown larvae raised on AZT. In contrast, knockdown of *wnd* showed an increase in the fragmentation of terminal dendrites, which was not significantly different from that induced by AZT in WT larvae (Fig. 4A, B, E). To further confirm these observations, we performed the same experiment on *dSarm* mutants. SARM (**s**terile α-motif-containing and **arm**adillo-motif containing protein) also works in the Wallerian degeneration pathway and mutations in d*SARM* also suppress Wallerian degeneration ^70,71^. We found that mutations in *dsarm* were also ineffective in suppressing the fragmentation induced by AZT (Fig. 5 A, D, E). These data indicate that unlike paclitaxel-induced fragmentation, the mechanisms that underlie the fragmentation of terminal dendrites in larvae exposed to AZT are independent of the *wnd*/dlk and SARM pathway.

**Figure 4:**
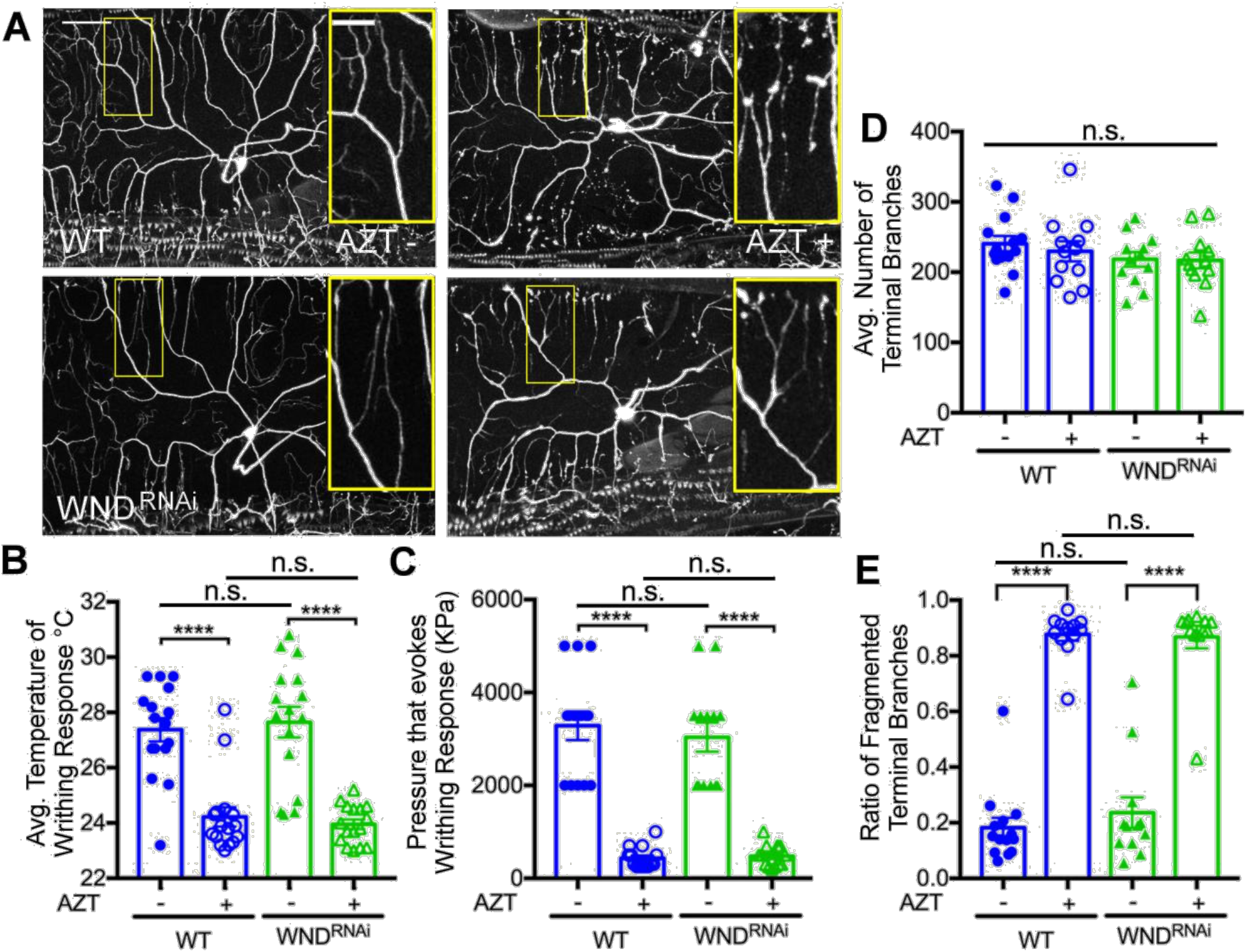
NRTI induced fragmentation of terminal dendrites is independent of the *wnd*/DLK pathway. **A**) Representative images of the C4da sensory neurons of third instar larvae showing Untreated WT (AZT−) and *wnd*^RNAi^ larvae (*wnd*^RNAi^, Bottom Left Panel) and AZT treated WT (AZT+), and *wnd*^RNAi^ (*wnd*^RNAi^ AZT+, Bottom right panel). **B**) Quantification of temperature required to elicit nociceptive behavior in AZT− and *wnd*^RNAi^ larvae. F(3, 60) = 25.0, p = 1.26E-10, 1-way ANOVA. Posthoc Bonferroni (AZT−, AZT+) p = 2.42E-6, (*wnd*^RNAi^, *wnd*^RNAi^ AZT+) p = 4.58E-7, (AZT−, *wnd*^RNAi^) p = 0.693, (AZT+, *wnd*^RNAi^ AZT+) p = 0.506. **C**) Quantification of pressure needed to induce mechanical nociception in AZT− and *wnd*^RNAi^ larvae. F(3, 50) = 48.9, p = 6.60E-15, 1-way ANOVA. Posthoc Bonferroni (AZT−, AZT+) p = 1.59E-9, (*wnd*^RNAi^, *wnd*^RNAi^ AZT+) p = 3.04E-8, (AZT−, *wnd*^RNAi^) p = 0.579, (AZT−, *wnd*^RNAi^ AZT+) p = 0.627. **D**) Quantification of terminal branches of C4da sensory neurons in AZT− and *wnd*^RNAi^ larvae. F(3, 46) = 0.930, p = 0.434, 1-way ANOVA. **E**) Quantification of proportion of C4da terminal branches that exhibit fragmentation induced by AZT in *wnd*^RNAi^ larvae. F(3, 46) = 92.8, p = 1.57E-19, 1-way ANOVA. Posthoc Bonferroni (WT, WT AZT) p = 3.82E-14, (*wnd*^RNAi^, *wnd*^RNAi^ AZT+) p = 4.74E-9, (AZT−, *wnd*^RNAi^) p = 0.405, (AZT+, *wnd*^RNAi^ AZT+) p = 0.844. WT=blue and *wnd*^RNAi^ larvae=green. Untreated (−); AZT (+). ****p < 0.0001; error bars = S.E.M.; scale bar = 50

**Figure 5:**
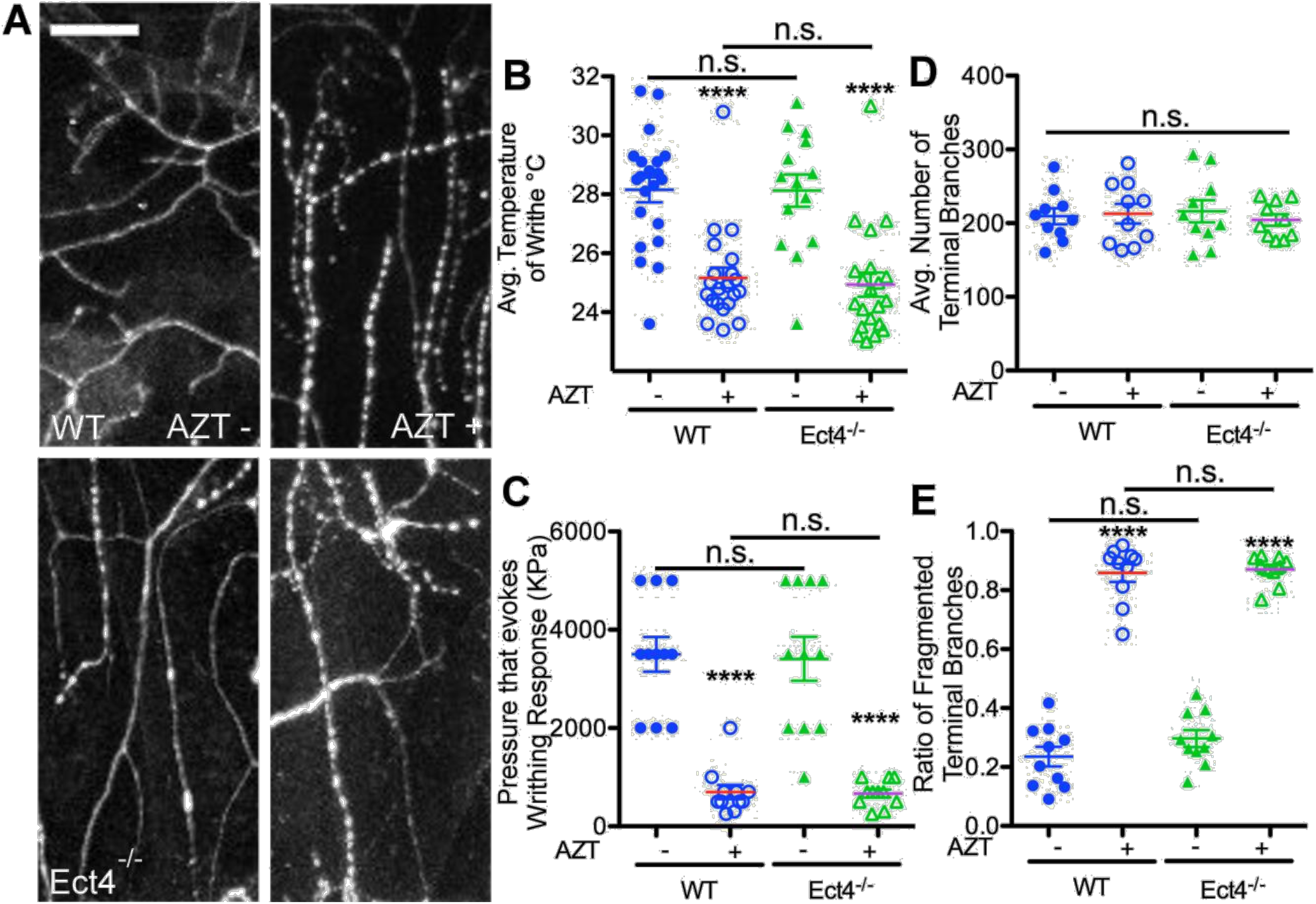
Pathways underlying NRTI induced degeneration are distinct from those regulating chemotherapy-induced degeneration. **A**) Representative images of the sensory neurons of third instar larvae comparing AZT− and AZT+ WT and Ect4^−/−^ larvae grown in AZT− and AZT+ food. **B**) AZT induces a significant decrease in the threshold temperature required to elicit the nociceptive behavior in both AZT+ and Ect4^−/−^ larvae. F(3, 93) = 20.1, p = 3.87E-10, 1-way ANOVA. Posthoc Bonferroni (AZT−, AZT+) p = 9.50E-8, (AZT-Ect4^−/−^, Ect4^−/−^ AZT+) p = 1.86E-5, (AZT−, Ect4^−/−^) p = 0.641, (AZT+, Ect4^−/−^ AZT+) p = 0.672. **C**) AZT induces a significant decrease in the threshold of mechanical stimulus required to elicit nociceptive behavior in both AZT− and AZT-Ect4^−/−^ larvae. F(3, 40) = 29.4, p = 3.34E-10, 1-way ANOVA. Posthoc Bonferroni (AZT−, AZT+) p = 3.81E-7, (AZT-Ect4^−/−^, Ect4^−/−^ AZT+) p = 6.56E-6, (AZT−, Ect4^−/−^) p = 0.874, (AZT+, Ect4^−/−^ AZT+) p = 0.871. **D**) The quantity of terminal branches of sensory neurons is unaffected by AZT in both WT AZT− and AZT-Ect4^−/−^ larvae. F(3, 36) = 0.173, p = 0.914, 1-way ANOVA. **E**) The proportion of terminal branches that exhibit fragmentation is increased by AZT exposure in both AZT− and Ect4^−/−^ larvae. AZT− quantification in blue and AZT-Ect4^−/−^ larvae quantification in green. F(3, 36) = 154, p = 1.32E-20, 1-way ANOVA. Posthoc Bonferroni (AZT−, AZT+) p = 5.61E-11, (AZT− Ect4^−/−^, Ect4^−/−^ AZT+) p = 7.90E-13, (AZT−, AZT− Ect4^−/−^) p = 0.182, (AZT+, Ect4^−/−^ AZT+) p = 0.724. Vehicle treated= AZT−; AZT treated= AZT+. N.S. = p > 0.05, ****p < 0.0001; error bars = SEM; scale bar = 20 μm.

Although *wnd*/DLK knockdown (or d*SARM* mutants) did not suppress the fragmentation of dendrites, we asked whether these pathways might still be able to suppress the nociceptive hypersensitivity induced by the NRTIs without suppressing the fragmentation. However, knockdown of *wnd*/DLK (Fig. 4 B, C) or mutations in d*Sarm* (Fig. 5 B, C) did not suppress the nociceptive hypersensitivity induced by AZT, indicating that these pathways may not play a significant role in suppressing the neurotoxicity induced by NRTIs. Finally, we also performed general motility tests on d*Sarm* and *wnd*^RNAi^ lines and did not find any significant defects in the general motility of these larvae (data not shown).

### Exposure to AZT leads to an increase in dynamic dendrites

To understand the cellular mechanisms that may underlie the terminal dendrite fragmentation, we turned to time-lapse imaging studies. Previous studies have suggested that chronic instability of dendrites might be a precursor for neurodegeneration (reviewed in ^84^). Therefore, we wanted to test whether the terminal dendrites that showed the fragmentation were more dynamic in AZT exposed larvae. To test this, we performed time-lapse imaging on C4da neurons labeled by GFP driven by ppk-Gal4^63^ on live animals without anesthesia (see materials and methods for details). Images were acquired from the entire dendritic field of one to two C4da neurons in between body segments 2-5, every 10 minutes for 3 hours. Random section of the image from each time point was compared to its previous time point to determine the number of branches that were extending, retracting, or sprouting. These analyses should reveal whether there is any difference between the dynamic nature of the dendrites between the two experimental conditions (AZT+ and AZT−). These analyses revealed that the number of terminal dendrites that were exhibiting dynamic changes was significantly increased in larvae raised on AZT (Fig. 6, also see Supplementary Movies 3[AZT−], 4 [AZT+]). These data suggest that WT larvae that were not raised on AZT have more stable terminal dendrites than those exposed to AZT. Furthermore, larvae whose dendrites were exposed to AZT first extended and then retracted the dendrites similar to a smaller percentage of WT dendrites not exposed to AZT (Fig. 6 and Supplementary movie 3). In contrast, majority of the dendrites exposed to paclitaxel showed only retractions and had significantly fewer extending or sprouting dendrites (Supplementary Fig. 7), further highlighting the possible underlying differences in the cellular mechanisms of fragmentation induced by AZT and paclitaxel. Finally, the live imaging experiments also showed gradual fragmentation of dendrites with time in AZT+ larvae (Supplementary Fig. 8), showing that the fragmentation observed in the dissected larvae was also apparent in live imaging. Based on these data, we conclude that exposure to AZT leads to more dynamic dendrites, which may make them vulnerable to fragmentation^84^.

**Figure 6:**
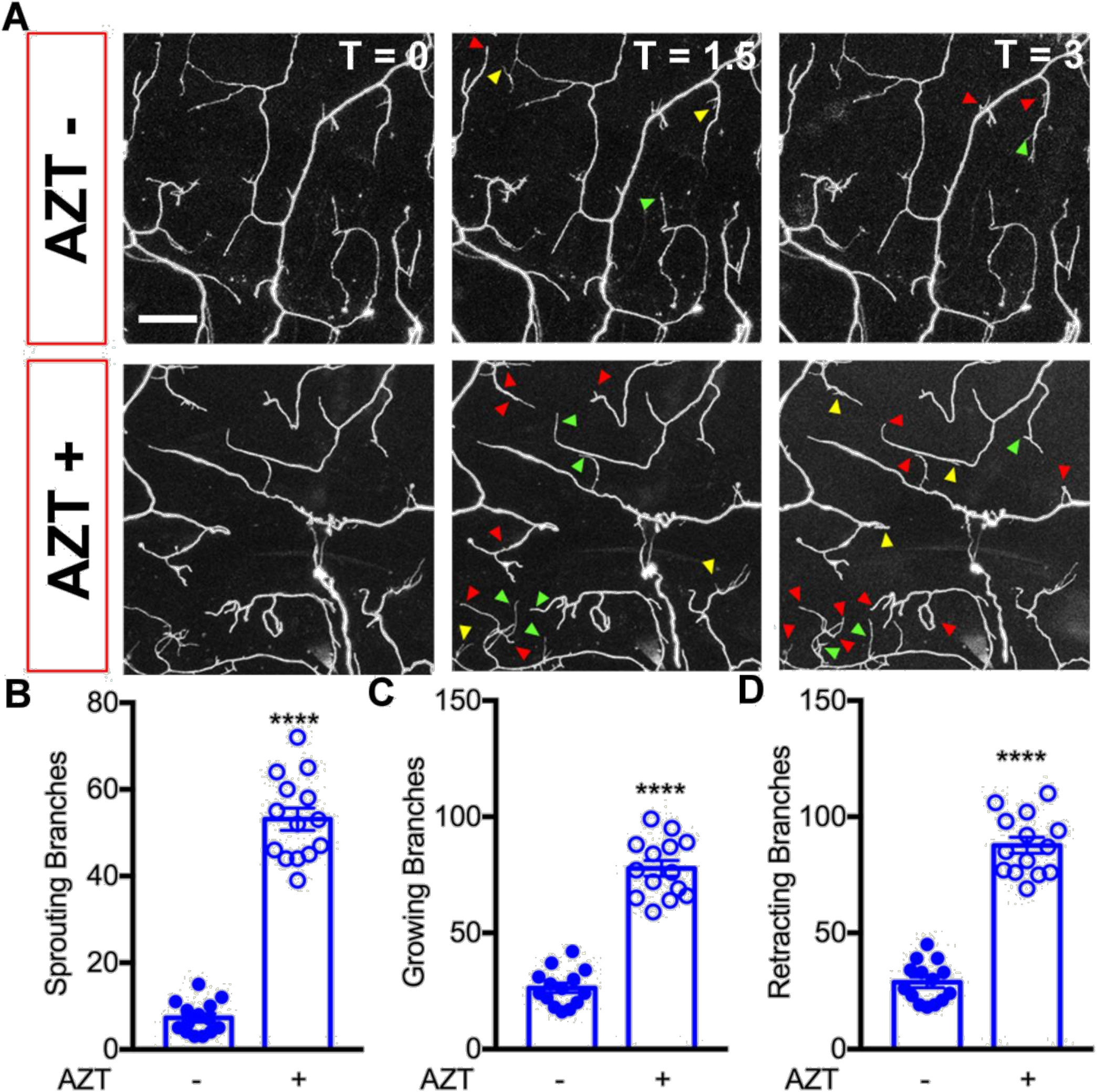
NRTI exposure increases the number of dynamic dendrites. **A**) Representative images of C4da terminal dendrites imaged over 3 hours in larvae raised on AZT− or AZT+ food. The time points shown are 0, 1.5 hours, and 3 hours. B-D) Quantification of terminal dendrites showing sprouting (**B**) (p = 2.34E-15, t-test), growing (**C**) (p = 3.77E-13, t-test), and retraction (**D**) p = 8.79E-14, t-test. AZT−; AZT+. ***p < 0.001; error bars = SEM, scale bar = 20 μm. Red arrowheads show retracting dendrites, Green arrow shows growing dendrites and yellow arrows show sprouting dendrites. Refer to Movie 3, 4 for video examples of live imaging frames and dendrite changes. Refer to supplementary Fig. 7 for Taxol effects on sensory neuron dynamics.

### Genetic restoration of dendritic stability suppresses AZT induced fragmentation and nociception

Because time-lapse imaging revealed that terminal dendrites of neurons exposed to AZT were more dynamic, we wondered whether the instability of dendrites might contribute to their fragmentation. To test this hypothesis, we genetically manipulated the stability of dendrites by modulating the levels of Par-1 kinase, a microtubule-associated serine-threonine kinase whose levels are important in regulating dendritic stability during development ^72^. During normal development of dendrites, an increase in Par-1 kinase leads to a decreased stability of dendrites and *vice versa*. Consistent with these findings, we found that increasing the levels of Par-1 kinase in the sensory neurons of WT larvae raised in AZT− food led to a significant decrease in the length of terminal dendritic branches, suggesting that they were unstable (Fig. 7). Decreasing the levels of Par-1 in larvae raised in AZT− food did not show any significant changes in dendrite morphology (Fig. 7). Consistent with our hypothesis, Par-1 knockdown larvae raised on AZT had significant suppression of terminal dendrite fragmentation (Fig. 7E). These data demonstrate that restoring the dendritic stability can suppress their fragmentation in flies raised on AZT.

**Figure 7:**
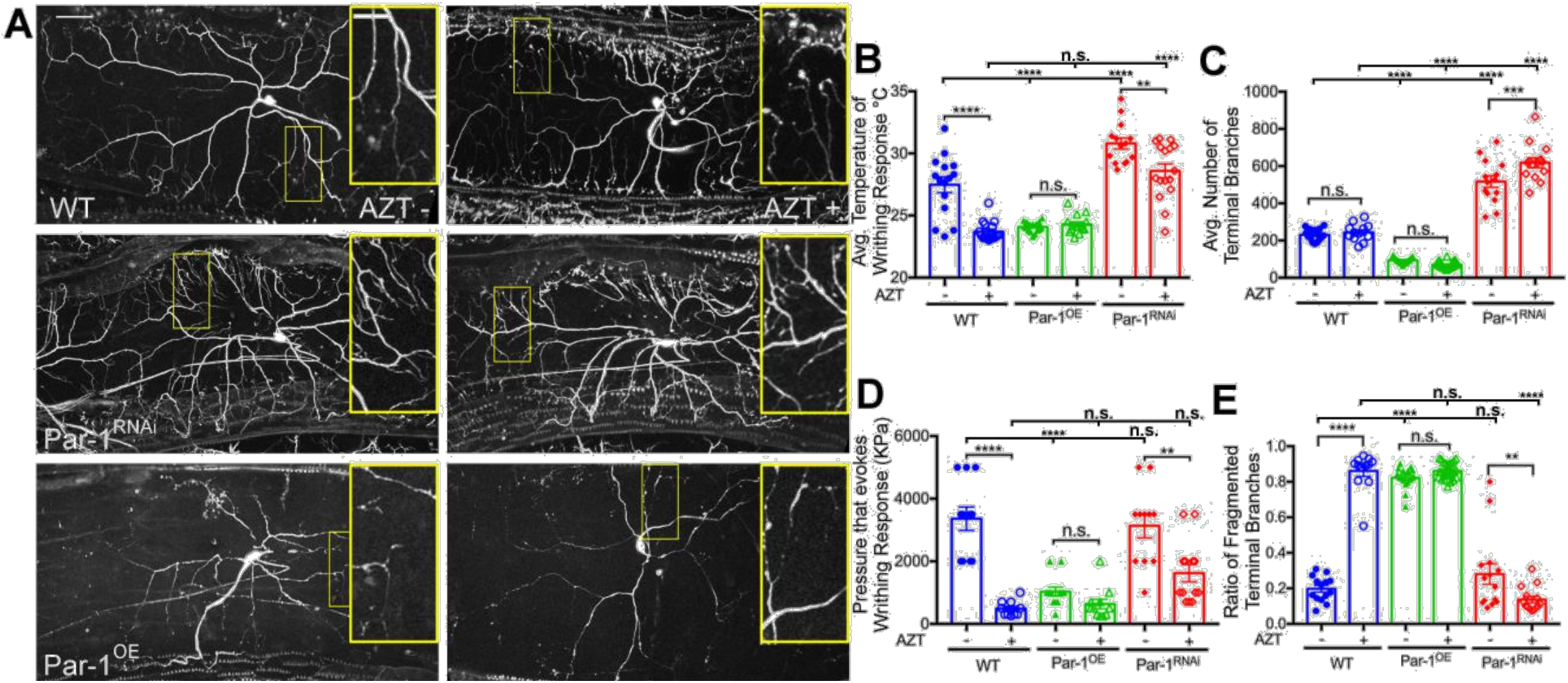
Restoration of dendrite stability suppresses fragmentation induced by AZT. **A**) Representative images of GFP labeled-C4da sensory neurons of third instar larvae raised on food containing vehicle (AZT−) or AZT (AZT+). The genotypes are indicated on the figure and are: AZT−, Par-1 overexpression, AZT− (Par-1^OE^), and AZT+ Par-1^RNAi^. **B**) Quantification of nociceptive response to thermal stimulus from AZT−, Par-1 overexpression (Par-1^OE^) and Par-1 knockdown (Par-1^RNAi^) lines on AZT− or AZT+ media. F(5, 85) = 49.0, p = 1.27E-23, 1-way ANOVA. Posthoc Bonferroni (AZT−, AZT+) p = 2.61E-6, (Par-1^OE^, Par-1^OE^ AZT+) p = 0.353, (Par-1^RNAi^, Par-1^RNAi^ AZT+) p = 0.00577, (WT AZT−, Par-1^OE^) p = 7.55E-6, (WT, Par-1^RNAi^) p = 2.53E-5, (WT AZT+, Par-1^OE^ AZT+) p = 0.0959, (WT AZT+, Par-1^OE^ AZT+) p = 4.49E-9. **C**) Quantification of number of terminal branches in the same genotypes as in B when raised on AZT− or AZT+ containing food. F(5, 90) = 206, p =1.02E-47, 1-way ANOVA. Posthoc Bonferroni (AZT−, AZT+) p = 0.383, (Par-1^OE^, Par-1^OE^ AZT+) p = 0.0870, (Par-1^RNAi^, Par-1^RNAi^ AZT+) p = 0.000527, (WT AZT−, Par-1^OE^) p = 2.98E-15, (WT AZT−, Par-1^RNAi^) p = 2.76E-9, (AZT+, Par-1^OE^ AZT+) p = 3.72E-18, (AZT+, Par-1^OE^ AZT+) p = 1.53E-11. **D**) Quantification of nociceptive response to mechanical stimulus of the same genotypes as in (B, C) when raised on AZT− or AZT+ food. F(5, 62) = 21.6, p = 1.87E-12, 1-way ANOVA. Posthoc Bonferroni (WT AZT−, WT AZT+) p = 8.39E-7, (Par-1^OE^, Par-1^OE^ AZT+) p = 0.0729, (Par-1^RNAi^, Par-1^RNAi^ AZT+) p = 0.00336, (WT AZT−, Par-1^OE^) p = 6.16E-6, (WT AZT−, Par-1^RNAi^) p = 0.676, (WT AZT+, Par-1^OE^ AZT+) p = 0.404, (WT AZT+, Par-1^OE^ AZT) p = 0.0583. **E**) Quantification of terminal branch fragmentation in the same genotypes as in (B-D) in AZT− or AZT+ food. F(5, 90) = 184, p = 1.05E-45, 1-way ANOVA. Posthoc Bonferroni (WT AZT−, WT AZT+) p = 7.16E-16, (Par-1^OE^, Par-1^OE^ AZT+) p = 0.0780, (Par-1^RNAi^, Par-1^RNAi^ AZT+) p = 0.00265, (WT AZT−, Par-1^OE^) p = 6.02E-22, (WT AZT−, Par-1^RNAi^) p = 0.188, (WT AZT+, Par-1^OE^ AZT+) p = 0.967, (WT AZT+, Par-1^OE^ AZT+) p = 8.00E-17. WT=blue, Par-1^OE^=green, and Par-1^RNAi^ =red. AZT−; AZT+. **p < 0.01, ***p < 0.001, ****p < 0.001; error bars = SEM, scale bar = 50 μm; inset scale bar = 20 μm.

Next, we asked whether restoring dendritic stability could also suppress the nociceptive hypersensitivity induced by AZT. To test this, we raised WT and WT larvae expressing Par-1^RNAi^ specifically in their sensory neurons ^73,74^ on AZT and subjected them to both thermal and mechanical nociception paradigms. Indeed, we observed that restoring dendritic stability led to significant suppression of both thermal as well as mechanical nociceptive hypersensitivity (Fig. 7B, D). These data strongly suggest that alteration in dendritic stability may contribute to the sensory neuron degeneration and degeneration of the sensory neurons is likely to cause nociceptive hypersensitivity in the *Drosophila* model of NRTI induced neurotoxicity.

## Discussion

In this study, we show that *Drosophila* can be used as a model organism for investigating the mechanisms underlying peripheral sensory neuropathy and nociceptive hypersensitivity induced by NRTIs. Furthermore, our data suggest that NRTIs affect the stability of sensory neurons, which may make them susceptible to degeneration ^75,76,^ and together, these may contribute toward the development of nociceptive hypersensitivity induced by the NRTIs in this model. While this model is an important first step toward understanding the neurotoxicity of NRTIs, further work needs to be done to establish whether these changes are observed in vertebrate models, which we are currently investigating and unpublished data from our labs supports that conclusion (Bush, Tang, and Wairkar, unpublished data). Finally, further research on this topic is necessary because pain hypersensitivity has been linked to the development of chronic pain ^77^, a symptom prevalent in patients living with HIV^78–80^.

### *Drosophila* as a model to understand the mechanisms of NRTI-induced PSN

*Drosophila* has proved to be an excellent model in understanding the mechanisms of nociception ^33,37,44,45^. The characteristic “corkscrew” like behavior of *Drosophila* larvae to noxious stimuli has been exploited to screen for proteins that are required for nociception^52^. These initial studies have led the way to recent studies that have been instrumental in understanding the underlying mechanisms of peripheral neuropathy caused by the use of chemotherapeutic drugs such as Taxol (Paclitaxel) and vincristine ^41,49^. Our study has furthered the use of *Drosophila* as a model to understand the mechanisms of nociceptive hypersensitivity induced by NRTIs, opening up the possibility of performing genome-wide forward genetic screens to elucidate the detailed mechanisms of NRTI induced neurotoxicity. Since many of the components for sensation, regulation, and integration of nociceptive signals are conserved from flies to vertebrates and humans ^32^, this model may prove to be useful in shedding light onto the mechanisms of neurotoxicity induced by NRTIs. Finally, many signaling mechanisms underlying nociception are also conserved in *Drosophila* ^37,81^. Given the easy access to the peripheral nervous system ^56^ and the lower costs of performing unbiased forward genetic screens together with a plethora of genetic tools makes *Drosophila* a powerful model to investigate the mechanisms underlying neurotoxicity of NRTIs and possibly, by the other components of anti-retroviral drugs such as the protease inhibitors.

### Stability of synapses and nociception

It is generally believed that activity patterns regulate the stability of synaptic communications. For example, during development, circuit refinement occurs based on activity ^82,83^ and active synapses are usually the ones that are stabilized. These connections can last for a long time, sometimes for decades in humans ^84^. Indeed, loss of spine and dendrite stability has been observed in some psychiatric disorders and neurodegenerative disorders like Alzheimer’s disease ^84^. A recent study in a mouse model of Tauopathy also suggests that loss of synapse stability might precede the overt loss of neurons in these neurodegenerative diseases ^85^. Consistent with these hypotheses, we found that exposure to NRTIs causes a loss of sensory neuron stability. Importantly, restoring the stability to the neurons led to the suppression of NRTI-induced sensory neuron fragmentation as well as nociception. These data suggest that synapse instability might underlie the PSN found in patients on chronic ART therapy and that restoring the stability of peripheral neurons could be protective. However, this idea needs to be further tested in vertebrate models. Also, more work needs to be done to understand the nature of this instability and the pathways that regulate it and ultimately, whether the instability of dendrites drives the hypersensitivity in pain underlying the use of ART and perhaps, in other chronic pain conditions. The combination of forward genetic screens in *Drosophila* and robust rodent pain models might be an effective way to address these questions in the future.

### Chemotherapy-induced PSN versus NRTI-induced PSN

Chemotherapy-induced peripheral neuropathy (CIPN) is a significant problem associated with the use of chemotherapy drugs to treat cancer patients ^88^. The symptoms associated with CIPN present similar to that induced by NRTIs i.e., PSN. Recently, there has been a flurry of animal models of CIPN both in vertebrates and in flies ^41,49,89–91^. These studies on CIPN in model systems suggest that signaling mechanisms that play a vital role in axon degeneration ^41,49^ (which undergoes a characteristic degeneration associated with axonal injury^67^), might also underlie CIPN. One of the common pathways associated with both axon degeneration and CIPN is the Di-Leucine zipper kinase (Dlk)-pathway. Reducing the levels of *dlk* (*wnd* in flies^92^) results in the protection of distal axons and, it provides a similar level of protection in CIPN in *Drosophila* models ^41,49,93^. Interestingly, however, our data show that the NRTI-induced fragmentation of peripheral sensory neurons is not suppressed by reducing the levels of *wnd* (Dlk), suggesting that the pathways that underlie CIPN and NRTI-induced fragmentation of dendrites may be different. Intriguingly, a recent model of CIPN that uses more optimized doses of paclitaxel suggests that dendrite stability may also play a role in the degeneration observed in paclitaxel-induced neuropathy models ^41^. Together, these data suggest that even if there were a divergence in signaling pathways that cause PSN, some of the mechanisms that lead to the sensory neuron degeneration might be common and may hinge on regulating the stability of peripheral sensory neurons.

## Data Availability Statement

The authors confirm that the data supporting the findings of this study are available within the article [and/or] its Extended Data Figures.

## Acknowledgments

We are grateful to the members of Michael Galko lab and the Tang lab for the design of Von Frey filaments. We are indebted to Dr. Michael Galko and his team especially, Yan Wang for helping us perform the heat probe nociception assays. We are also thankful to Ms. Bianca Gozales for help with the project. Keegan Bush was supported by a fellowship from the neuroscience and cell biology department of UTMB and Mitchell center for neurodegenerative diseases. We are also thankful for funding from the NINDS R56NS105681, Alzheimer’s Association NTF award, and STARs award from the UT system to YPW. SJT was supported by NIH grants: R01NS095747, R01NS079166, R01DA036165. Finally, we would like to thank Dr. Raji Natarajan for scientific edits. The authors declare no conflicts of interest.

## Author Contributions

KMB, KRB, JAM, performed research

KMB, SJT, and YPW designed research

KMB and YPW analyzed data.

KMB and YPW wrote the paper

## Supplementary Movies and Methods

**Movies 1 and 2: Representative larval behavior in thermal nociception assay.**

**Movie 1**) Video showing normal larval behavior at sub-nociceptive temperature and characteristic nociceptive larval writhe at noxious temperatures. **Movie 2**) Video shows behavior of larvae raised on normal (WT-Left), and AZT (right) containing food. Frames are time synced from the same video. The temperature frame includes the output of the thermal probe inserted into one of the wells filled with equal amounts of PBS as the one with larvae for accurate temperature measurement.

**Movies 3 and 4: Example of live imaging of distal dendrites in AZT− and AZT+ conditions.**

Video showing 50μm by 50μm zoom of representative live imaging of larval terminal dendrites. Live imaging of larva grown in vehicle (**Movie 3**) or AZT (**Movie 4**) food show retraction (red arrowheads), elongation (Green arrowheads) and sprouting (Yellow arrowheads).

### Supplementary Data Methods

#### *Drosophila* NRTI treatment for newer NRTIs

All flies were raised at room temperature (22°C) with natural day night cycle. WT (*Canton S*) flies were placed in vials containing 2.5ml of instant *Drosophila* media (“blue food”, Carolina Biological Supply, Burlington, NC) made with water base treated with DMSO (Vehicle), 30μM Taxol (Paclitaxel, Sigma, St. Louis, MO) in DMSO, AZT 0.26μg/mL (main concentration used throughout except in the dose response experiments), Dose response experiments: AZT: 0.026 μg/mL, 0.13 μg/mL, 0.26 μg/mL, 1.3 μg/mL, 2.6 μg/mL, 13 μg/mL), ddC (0.0014 μg/mL, 0.014 μg/mL, 0.07 μg/mL, 0.14 μg/mL, 0.28 μg/mL, 0.56 μg/mL, 0.84 μg/mL, 1.12 μg/mL, 1.4 μg/mL, or 2.8 μg/mL, Abacavir (ABC) (lot 11 ND21-134-4): 0.026 μg/mL, 0.13 μg/mL, 0.26 μg/mL, 1.3 μg/mL, 2.6 μg/mL, 13 μg/mL, 26 μg/mL, Emtricitabine (FTC) (lot11 NB16-106-3): 0.0087 μg/mL, 0.043 μg/mL, 0.087 μg/mL, 0.43 μg/mL, 0.87 μg/mL, 4.3 μg/mL, 8.7 μg/mL, Tenovir disoproxil fumarate (Tenovir) (lot 20 NG41-110-2) 0.013 μg/mL, 0.065 μg/mL, 0.13 μg/mL, 0.65 μg/mL, 1.3 μg/mL, 6.5 μg/mL, 13 μg/mL).A stock of 1 mg/ml paclitaxel in DMSO was utilized for taxol containing food preparations with equivalent DMSO controls. All the NRTIs were obtained from the NIH AIDS Reagent program (http://aidsreagent.org).

**Supplementary Figure 1:**
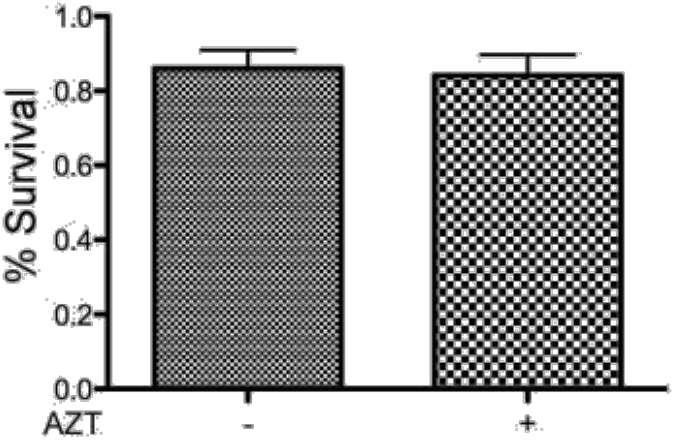
Effects of AZT on larval survival. Quantification of larval survival in vehicle and AZT food. 50 embryos of WT *Drosophila* were transferred to AZT− and AZT+ food. There was no significant difference in the number of larva and/or adult flies that survived. p = 0.782, t-test.

**Supplementary Figure 2:**
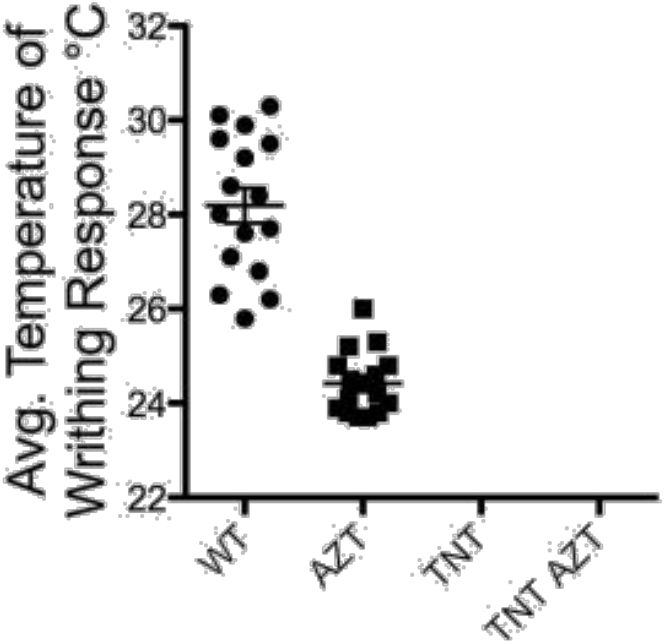
Nociceptive effects of NRTI are dependent on C4da sensory neurons. Quantification of temperatures required for the writhing response in larvae raised on AZT− and AZT+ food with and without driving tetanus toxin light chain (TNT) in the C4da neurons. The larvae were non-responsive to changes in temperature when tetnus toxin light chain was expressed in the C4da sensory neurons (last two bars). (WT, AZT) p=3.00E-9, t-test. TNT larva were non-responsive to the thermal stimulus.

**Supplementary Figure 3:**
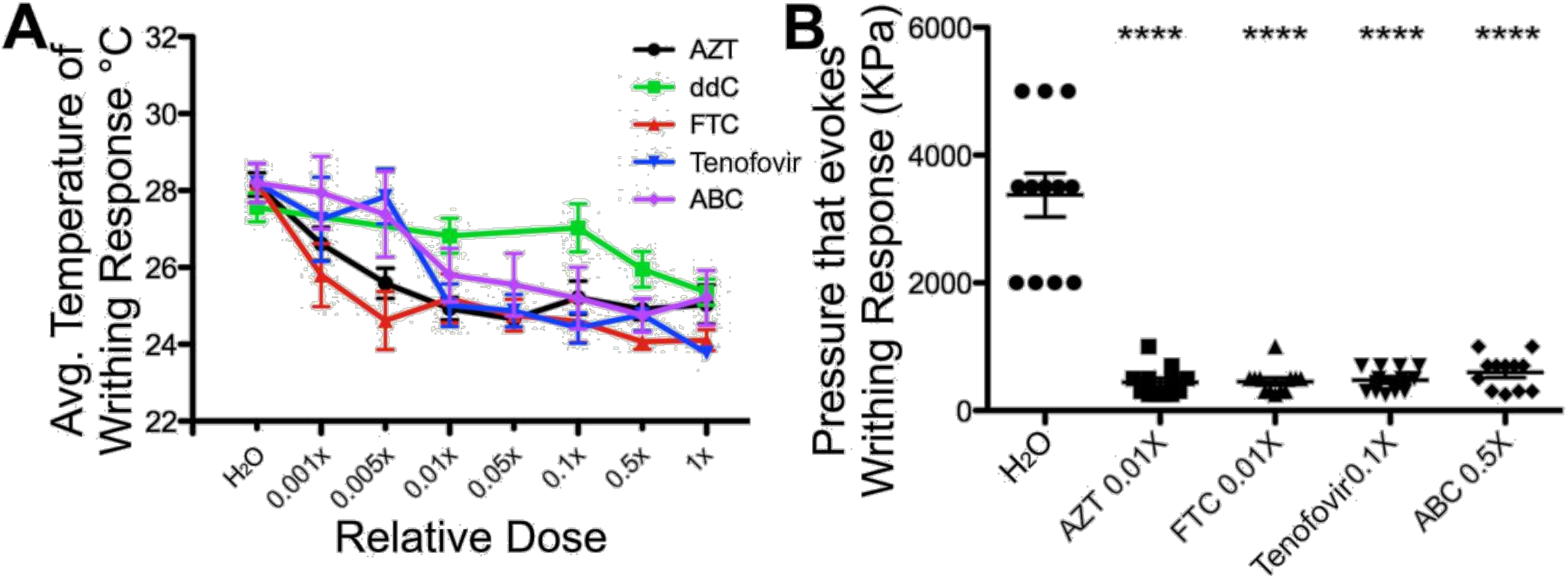
Newer NRTIs also induce thermal and nociceptive hypersensitivity in *Drosophila* model. **A)** Quantification of thermal response of larva to various (noted on figure) NRTIs. **B)** Quantification of mechanical hypersensitization of various NRTIs. F(4, 55) = 62.0, p = 9.95E-20, 1-way ANOVA. Posthoc Bonferroni (WT (Water), AZT+ 0.01X) p = 3.48E-6, (WT (Water), ABC+ 0.5X) p = 5.38E-6, (WT (Water), FTC+ 0.01X) p = 4.53E-6, (WT (Water), Tenofovir 0.1X) p = 3.75E-6. ****p < 0.0001; error bars = S.E.M.

**Supplementary Figure 4:**
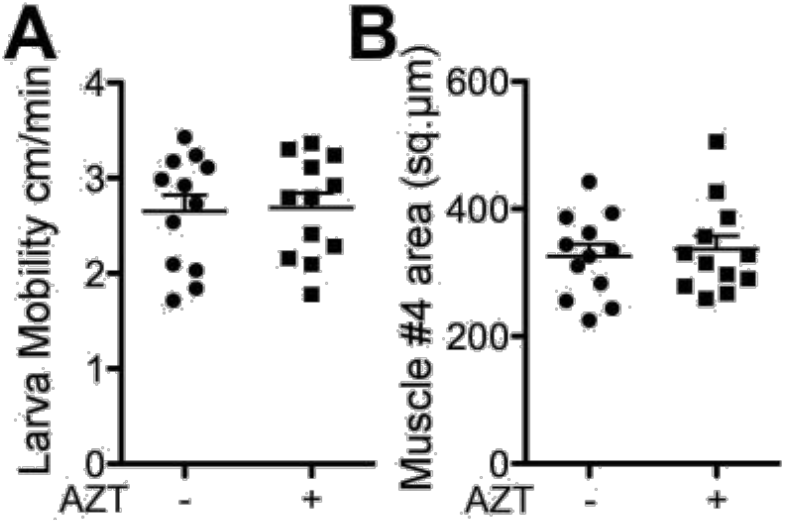
Larval motility and musculature are unaffected by AZT. **A)** Quantification of larval motility. There was no significant difference between larva raised on AZT− or AZT+ food. p = 0.873, t-test. **B**) Quantification of larva muscle 4 area (μm^2^) from abdominal segment 3 and 4. There was no significant difference in the muscle area of larvae raised on AZT− or AZT+ food. p = 0.966, t-test.

**Supplementary Figure 5:**
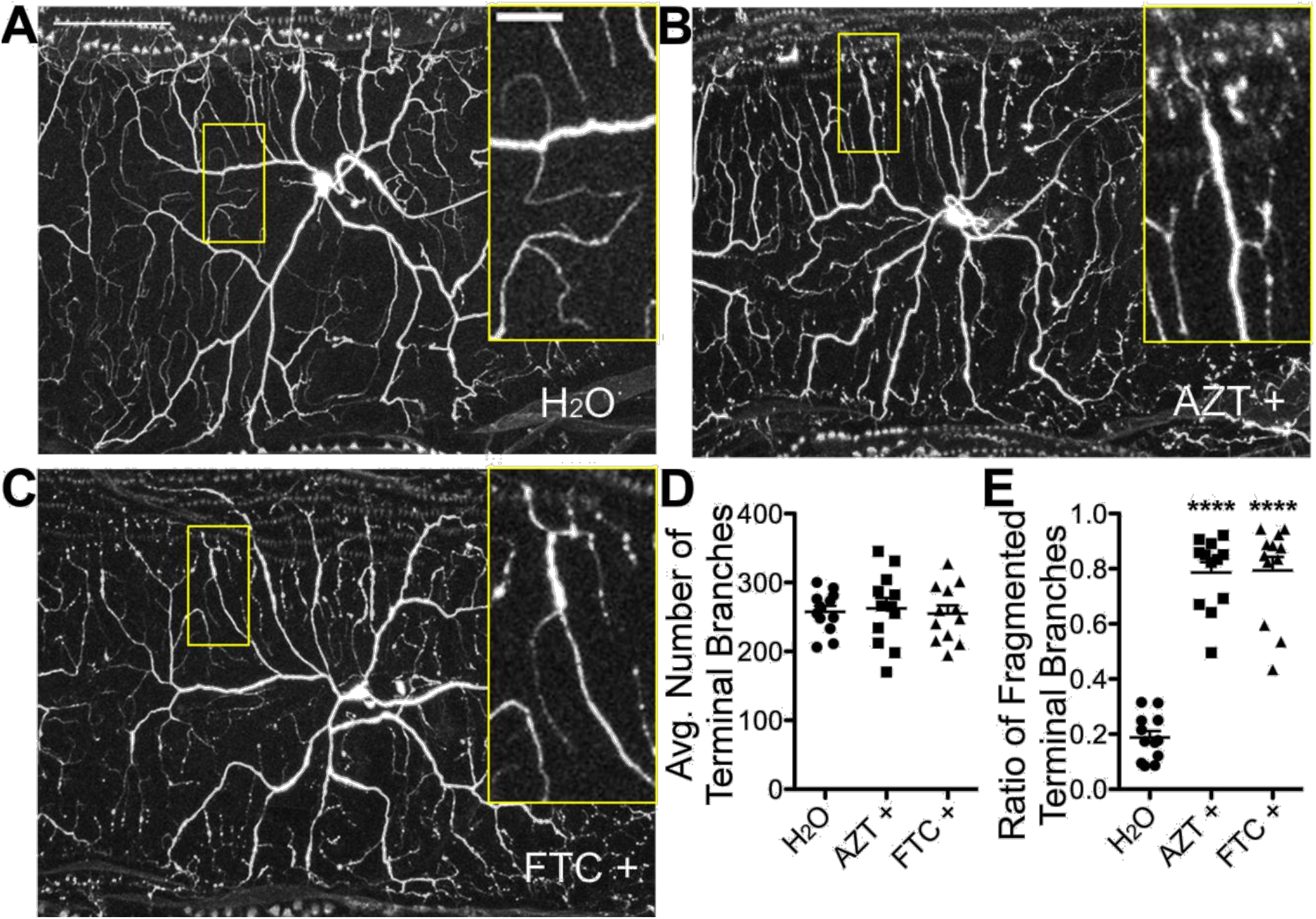
Exposure to FTC also leads to sensory neuron degeneration. **A)** Representative image of GFP labeled-C4da sensory neurons of third instar larvae raised on control food containing water (H_2_O). **B)** on food containing AZT and **C)** on food containing FTC. **D)** Quantification of terminal branches of C4da sensory neurons of larvae raised on control (H2o), AZT, and FTC. F(2, 33) = 0.0983, p = 0.907, 1-way ANOVA. **E)** Quantification of proportion of C4da terminal branches that exhibit fragmentation in identical genotypes as E. F(2, 33) = 81.2, p = 1.80E-13, 1-way ANOVA. Posthoc Bonferroni (WT, AZT 0.01X) p = 6.99E-8, (WT, FTC 0.01X) p = 5.75E-7. ****p < 0.0001; error bars = S.E.M.; scale bar = 50 μm; inset scale bar = 20 μm.

**Supplementary Figure 6:**
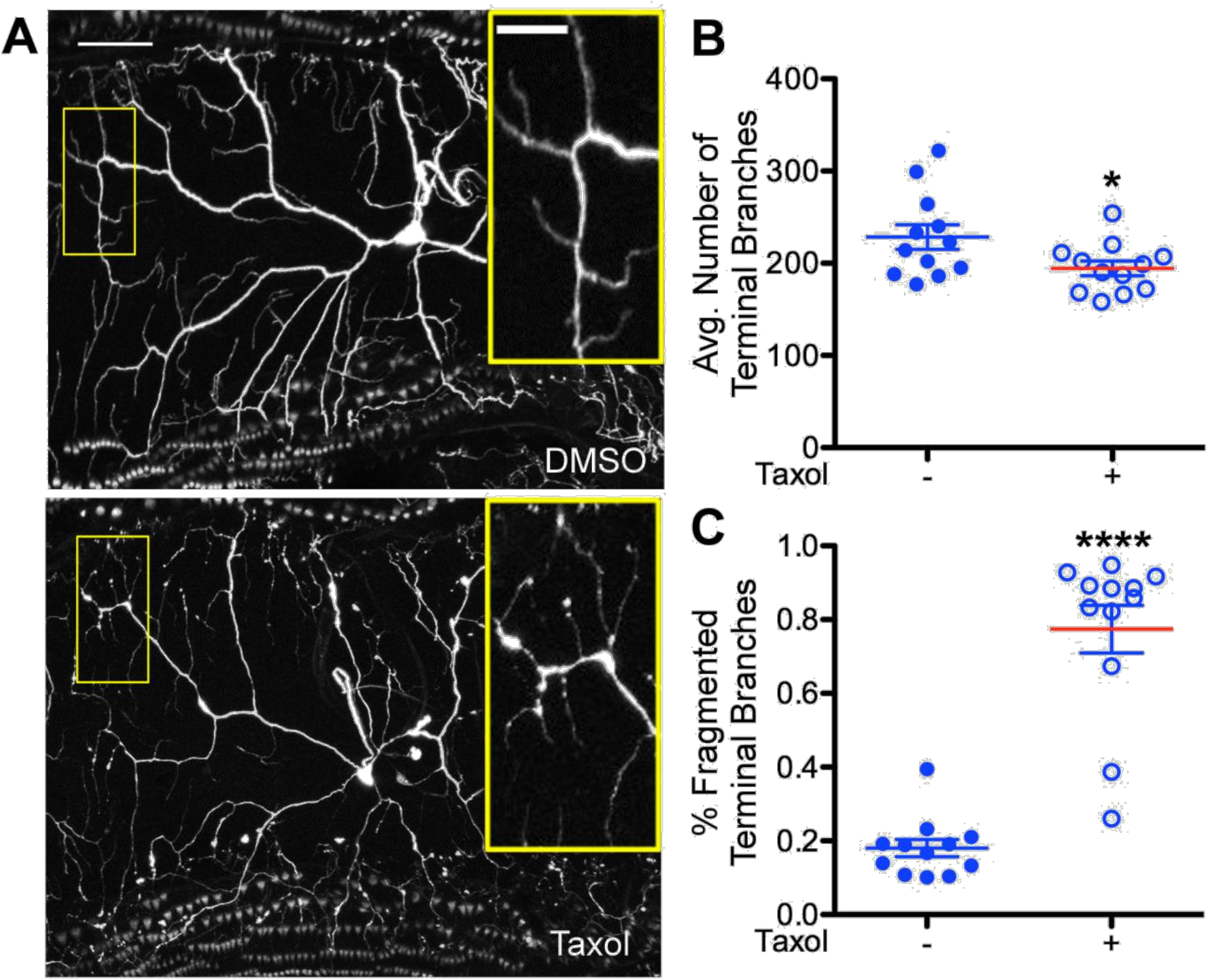
Taxol exposure also induces degeneration of sensory neurons. **A**) Representative images of the C4da sensory neurons of third instar larvae raised on vehicle (DMSO) and Taxol containing food. **B**) The number of terminal branches of C4da sensory neurons is decreased in larvae raised on Taxol. p = 0.0379, t-test. **C**) The proportion of C4da terminal branches that exhibit fragmentation is increased when the larvae are raised on Taxol containing food. p = 1.53E-8, t-test. Vehicle −; Taxol +. N.S. = p > 0.05, ****p < 0.0001; error bars = SEM; scale bar = 50 μm; inset scale bar = 20 μm.

**Supplementary Figure 7:**
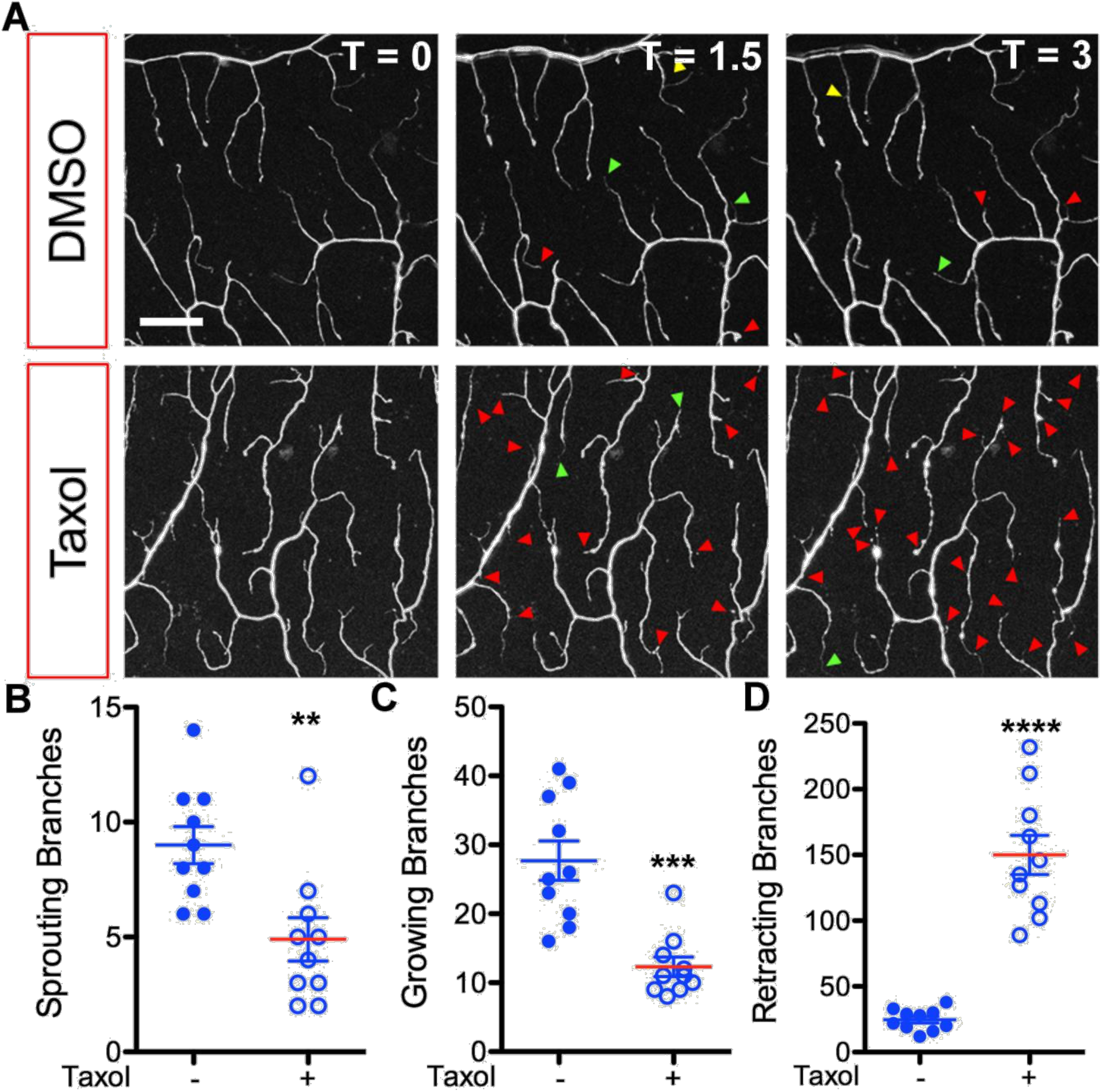
Taxol exposure induces increased dynamic changes in terminal dendrites of C4da sensory neurons. **A**) Representative images of C4da dendrites from third instar larvae raised on vehicle (DMSO) or Taxol containing food over a series of time periods (0, 1.5 hours, and 3 hours). **B**, **C**) The number of dendrites exhibiting dynamic changes in the form of growing (p = 0.00398, t-test), and sprouting (p = 0.000132, t-test) is decreased with exposure to Taxol while the quantity of retracting branches (**D**) is increased. p = 1.40E-7, t-test. Vehicle −; Taxol +. N.S. = p > 0.05, N.S. = p > 0.05, ***p < 0.001; error bars = SEM, scale bar = 20 μm.

**Supplementary Figure 8:**
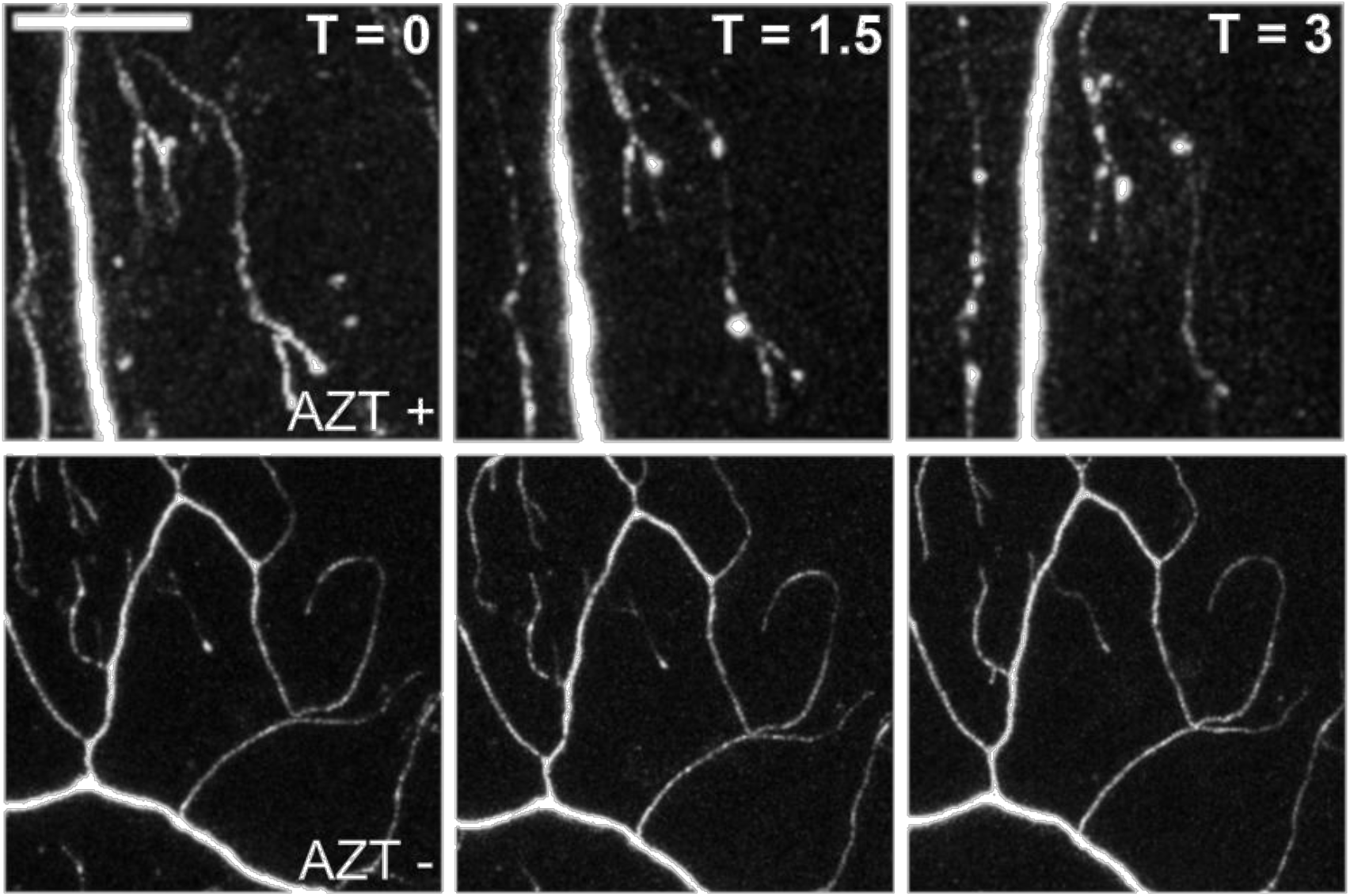
Example of dendrite fragmentation in AZT-Live Imaging. Representative images of fragmentation in C4da dendrites over time in larvae exposed to AZT and WT at time 0, 1.5 hours, and 3 hours. AZT top, WT bottom. Scale Bar = 20 μm.

